# Alignment of genetic differentiation across trophic levels in a fig community

**DOI:** 10.1101/2021.12.21.473707

**Authors:** Gavin C. Woodruff, John H. Willis, Patrick C. Phillips

## Abstract

Ecological interactions can generate close associations among species, which can in turn generate a high degree of overlap in their spatial distributions. Co-occurrence is likely to be particularly intense when species exhibit obligate comigration, in which they not only overlap in spatial distributions but also travel together from patch to patch. In theory, this pattern of ecological co-occurrence should leave a distinct signature in the pattern of genetic differentiation within and among species. Perhaps the most famous mutual co-isolation partners are fig trees and their co-evolved wasp pollinators. Here, we add another tropic level to this system by examining patterns of genomic diversity in the nematode *Caenorhabditis inopinata*, a close relative of the *C. elegans* model system that thrives in figs and obligately disperses on fig wasps. We performed RADseq on individual worms isolated from the field across three Okinawan island populations. The male/female *C. inopinata* is about five times more diverse than the hermaphroditic *C. elegans*, and polymorphism is enriched on chromosome arms relative to chromosome centers. *F*_*ST*_ is low among island population pairs, and clear population structure could not be easily detected among figs, trees, and islands, suggesting frequent migration of wasps between islands. Moreover, inbreeding coefficients are elevated in *C. inopinata*, consistent with field observations suggesting small *C. inopinata* founding populations in individual figs. These genetic patterns in *C. inopinata* overlap with those previously reported in its specific fig wasp vector and are consistent with *C. inopinata* population dynamics being driven by wasp dispersal. Thus, interspecific interactions can align patterns of genetic diversity across species separated by hundreds of millions of years of evolutionary divergence.

**Highlights:**

- The fig-dwelling female/male nematode *Caenorhabditis inopinata* is five times more diverse than its closest relative, the self-fertilizing nematode *C. elegans*.
- *C. inopinata* migrates frequently among three Okinawan islands despite high levels of inbreeding within individual figs.
- *C. inopinata* has patterns of genetic diversity that mirror its fig wasp vector.
- Ecological specialization aligns patterns of genetic differentiation in closely interacting species.

## Results and discussion

Transient resource patches are widespread and sustain myriad species across a range of spatial and temporal scales. For instance, whale falls emerge from ∼90 metric ton carcasses and can sustain dozens of metazoan species across decades [1, 2]. Conversely, decomposing fruit communities are orders of magnitude smaller in physical size but can likewise sustain complex communities of arthropods, mollusks, nematodes, yeast, and other microbes for weeks [3-5]. As such resources are temporary, most organisms adapted to such niches have dispersal mechanisms for finding new patches. These often involve interspecific interactions—phoresy is one such common ecological relationship wherein one species carries another, facilitating its dispersal [6]. Additionally, the carrier and the carried also frequently differ greatly in size: remora fish disperse on loggerhead sea turtles and whales [6, 7], and tardigrades disperse on birds [6, 8]. Such interactions require the comigration and colocalization of individuals of different species in space. Thus, interspecific relationships like phoresy have the potential to drive concordant patterns of spatial genetic differentiation among divergent species. That is, because they migrate and live together, it is reasonable to suspect that species in these kinds of relationships may have similar patterns of spatial genetic structure. Indeed, such patterns may not be limited to phoresy, but may also extend to other intimate interspecific relationships like those between hosts and parasites/pathogens [9]. How do such interactions affect patterns of genetic diversity across vast spatial scales, and does this lead to covariance of genetic diversity among community members inhabiting the same transient resource?

Here, we describe patterns of genomic diversity in the fig wasp-dispersed nematode, *Caenorhabditis inopinata*. Figs are ephemeral resources that harbor complex communities of wasps [10], ants [11], moths [12], mites [13], nematodes [14], and fungi [15]. Moreover, figs and their pollinating wasps represent a classic mutualism in ecology and evolution: figs need wasps for pollination, and wasps need figs for nutrients [16]. *C. inopinata*, a close relative of the *C. elegans* model system, proliferates in *Ficus septica* figs and disperses on its pollinating fig wasp, *Ceratosolen bisculatus* [17, 18]. Additionally, patterns of *C. bisculatus* genetic diversity throughout southeast Asia have been previously defined [19, 20]. Thus, this system is well-positioned to understand how interspecific interactions can drive genetic structure among community members of the same transient resource. We find that patterns of genomic diversity in *C. inopinata* reveal a striking similarity with those observed in its fig wasp vector across island population, suggesting intimate ecological interactions can drive the alignment of genetic differentiation across species separated by hundreds of millions of years of evolutionary divergence.

### *The genomic landscape of polymorphism in* C. inopinata

To understand patterns of genetic diversity in *C. inopinata*, we performed reduced-representation genome sequencing (i.e., bestRAD [21]) on twenty-four individual *C. inopinata* animals isolated from three islands of the Yaeyama archipelago, the westernmost islands of Okinawa, Japan (Fig 1; see Supplemental_tables.xlsx Sheets 1-2 for GPS coordinates and sample sizes). We also retrieved previously published alignment files of *C. elegans* wild isolates [22] to compare genome-wide levels of diversity between *C. inopinata* and *C. elegans*. Only alignments associated with the same restriction site we used for RAD in our *C. inopinata* samples were used for population estimates in *C. elegans*. After aligning and genotyping *C. inopinata* sequences, we retained 226,527 biallelic SNP’s across 77,737 loci of the *C. inopinata* genome (comprising 4,974,872 bp when including invariant sites; see Supplemental_tables.xlsx Sheets 3-5 for read counts as well as alignment and coverage information). These revealed even coverage across the genome after accounting for variation in restriction site density along *C. inopinata* chromosomes (Supplemental Figure 1). We then found that *C. inopinata* exhibits levels of genome-wide polymorphism that are about five times higher than *C. elegans* (Fig 2A; *C. inopinata*: mean π = 0.011, range = 0.0-0.085; *C. elegans* mean π = 0.0022, range = 0.0-0.060). Additionally, genome-wide polymorphism is three times more variable in *C. inopinata* than in *C. elegans* (*C. inopinata* π variance = 6.0 × 10^−5^; *C. elegans* π variance = 2.0 × 10^−5^). As previously reported in *C. elegans* and other *Caenorhabditis* species [23-25], polymorphism is elevated on chromosome arms relative to chromosome centers in *C. inopinata* (Fig. 2B; mean π_arms_ = 0.012, mean π_centers_ = 0.010). However, the magnitude of elevated diversity on chromosome arms is much lower in *C. inopinata* (Cohen’s *d* effect size = 0.26) than in *C. elegans* (Fig. 2B; Cohen’s *d* effect size = 0.61). Additionally, both *C. elegans* and *C. inopinata* reveal less enrichment of polymorphism on the arms of the X chromosome (Fig. 2A), although this difference is magnified in *C. elegans* (autosomes Cohen’s *d* effect size = 0.69; X Cohen’s *d* effect size = 0.083) compared to *C. inopinata* (autosomes Cohen’s *d* effect size = 0.27; X Cohen’s *d* effect size = 0.25). Taken together, this suggests that broad patterns of the genomic organization of polymorphism are conserved between these species despite differences in reproductive mode. But although *C. inopinata* has much higher levels of polymorphism than the hermaphroditic species *C. elegans*, this species is only modestly diverse compared to other outcrossing *Caenorhabditis* species (Supplemental Figure 2).

**Figure 1.**
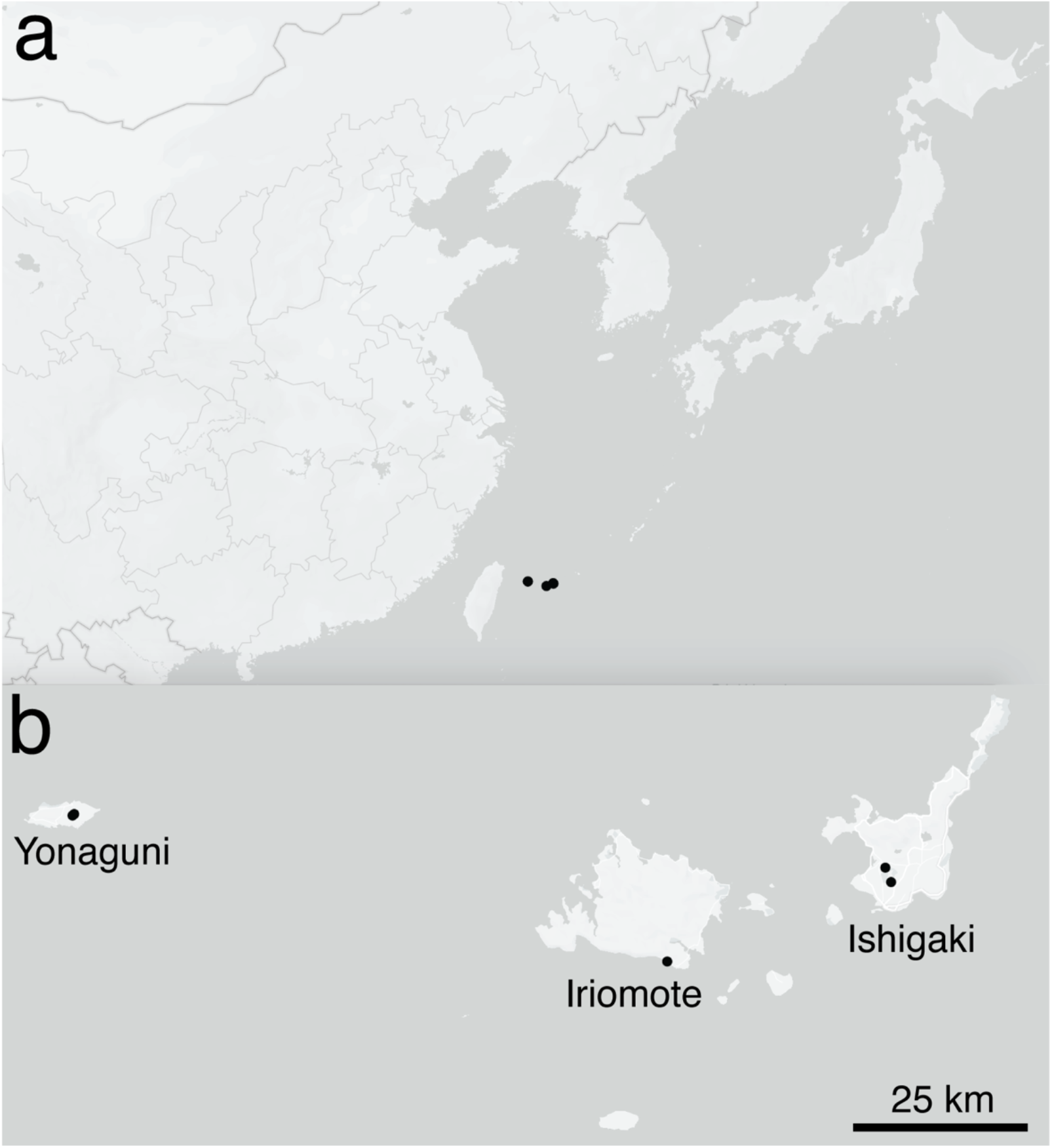
Geographic distribution of genotyped individuals. Twenty-four individual worms were genotyped from five trees across three islands in Okinawa. Maps were generated with Mapbox, OpenStreetMap, and their data sources. © Mapbox, © OpenStreetMap (https://www.mapbox.com/about/maps/; http://www.openstreetmap.org/copyright).

**Figure 2.**
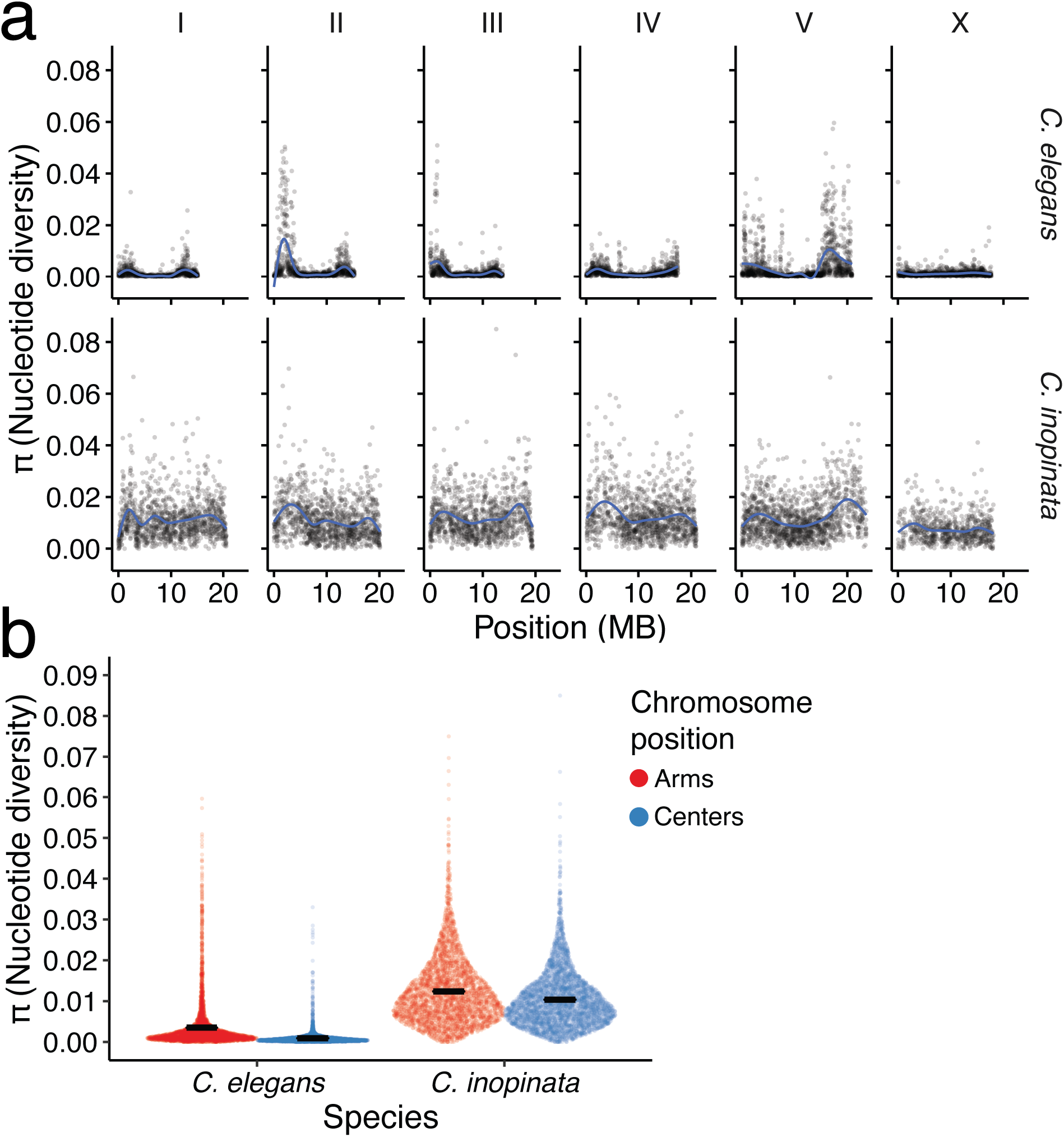
The genomic landscape of nucleotide diversity (π) in *C. inopinata* and *C. elegans*. (a) Nucleotide diversity across 10 kb genomic windows. Blue lines were fit by LOESS local regression. (b) Sina plots (strip charts with points taking the contours of a violin plot) revealing differences among chromosome arms and centers in *C. inopinata* and *C. elegans*. Horizontal black bars represent means. All *C. elegans* data were retrieved from the *CeNDR* project (https://www.elegansvariation.org/ ; [22]).

### C. inopinata *has elevated inbreeding and migrates frequently*

Because *C. inopinata* is distributed across the islands of Okinawa and Taiwan ([17, 18, 26]; and potentially throughout the islands of southeast Asia [19, 20, 27]), we explored the possibility of genetic differentiation among island populations. However, *F*_*ST*_ is generally low across all island population pairs (Fig 3; mean pairwise *F*_*ST*_ = 0.016; s.d. = 0.049; range = -0.44-0.75; Supplemental_tables.xlsx Sheet 6). Additionally, principal components analysis and network phylogenetic approaches do not reveal clusters or clades that correspond with island of origin (Supplemental Figure 3). Discriminant analysis of principal components [28] likewise does not reveal genetic partitioning across islands, plants, or figs across relevant values of *K* (Supplemental Figures 3-4). This all suggests that *C. inopinata* migrates frequently among the Yaeyama islands. At the same time, genome-wide estimates of *F*_*IS*_ (i.e., the inbreeding coefficient) are much higher and variable than those for *F*_*ST*_ (Figure 3; mean *F*_*IS*_ = 0.15, sd = 0.20, range= -0.73-1.0), suggestive of moderately high levels of mating among close relatives in *C. inopinata*. Thus these *F* statistics suggest the counterintuitive co-occurrence of frequent migration with high inbreeding in this species.

**Figure 3.**
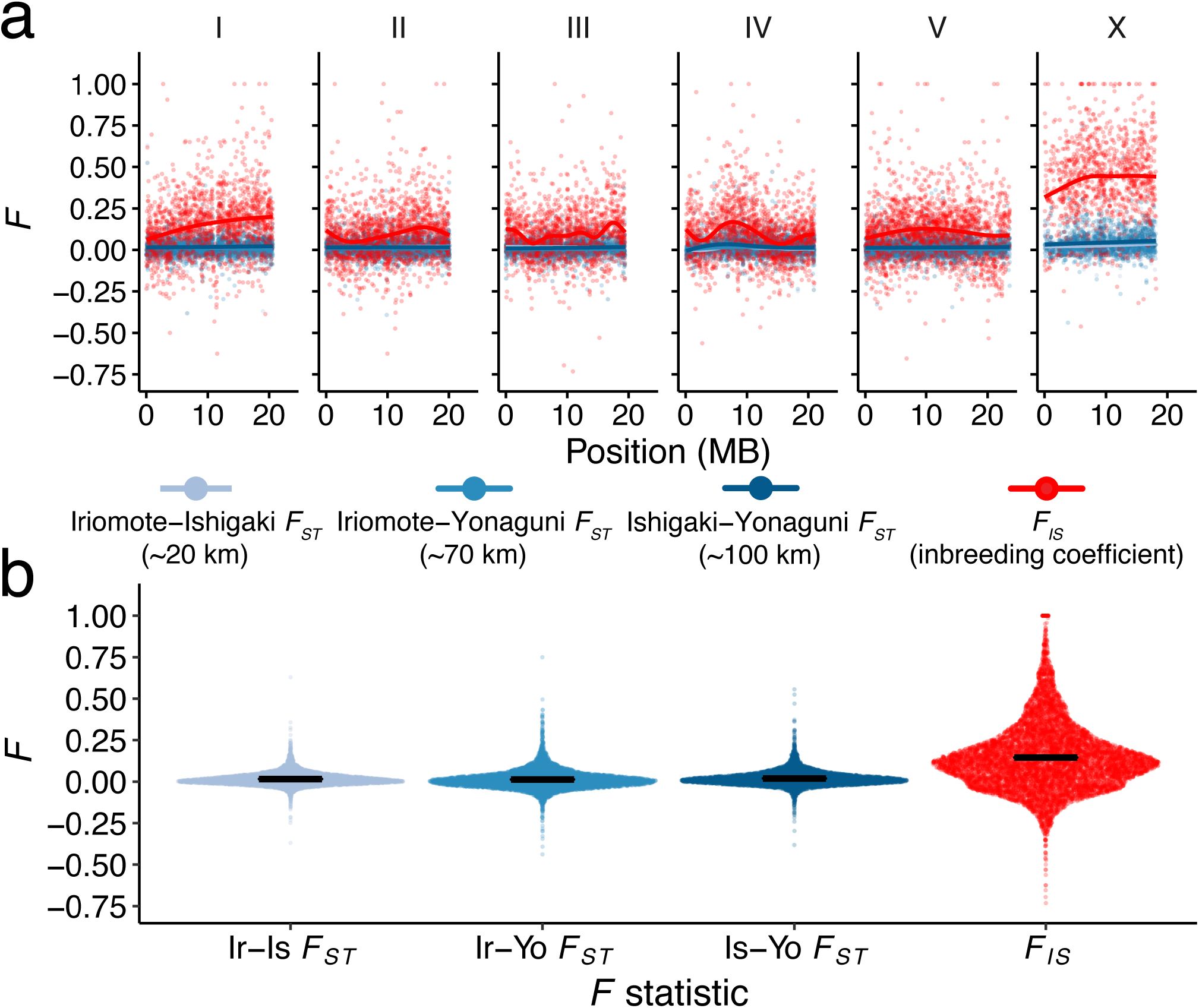
*C. inopinata* reveals low genetic differentiation among island populations despite high inbreeding. (a) Genomic landscapes of *F*_*IS*_ and island population pairwise *F*_*ST*_ across 10 kb genomic windows. Lines were fit by LOESS local regression. (b) Sina plots (strip charts with points taking the contours of a violin plot) revealing the distributions of the data in (a). Horizontal black bars represent means.

### C. inopinata *and its fig wasp carrier have comparable patterns of genetic diversity*

*C. inopinata* coexists with and is dispersed by the pollinating fig wasp *Certaosolen bisculatus* [17, 18]. We reasoned its patterns of genetic diversity across islands may be explained by its natural history and relationship with fig wasps. To address this, we retrieved previous *C. inopinata* ecological ([18]; Fig. 4A-B) and *C. bisculatus* population genetic ([19, 20]; Fig. 4C) data to situate the above results in their broader ecological context. Indeed, *F. septica* figs tend to be pollinated by a small number of pollinating wasps (Fig. 4A; mean = 1.9, range = 0-11; N = 162), and most wasps carry only a small number of *C. inopinata* animals (Fig. 4B; mean = 0.90, range = 0-6, N = 29) [18]. Moreover, two previous studies of *C. bisculatus* examined differentiation among populations that both included islands separated by open ocean in Indonesia [19] and Taiwan (Taiwan and Lanyu Island; [20]). Both studies revealed low differentiation among populations (mean pairwise *F*_*ST*_ in [19] = 0.015; mean pairwise *F*_*ST*_ in [20] = 0.0047). They also observed high inbreeding coefficients in *C. bisculatus* populations (mean *F*_*IS*_ [19] = 0.31; mean *F*_*IS*_ [20] = 0.24). Thus, *C. bisculatus* likewise likely migrates frequently across islands and exhibits high levels of inbreeding [19, 20], mirroring our observations of the worms they carry (Fig. 4C).

**Figure 4.**
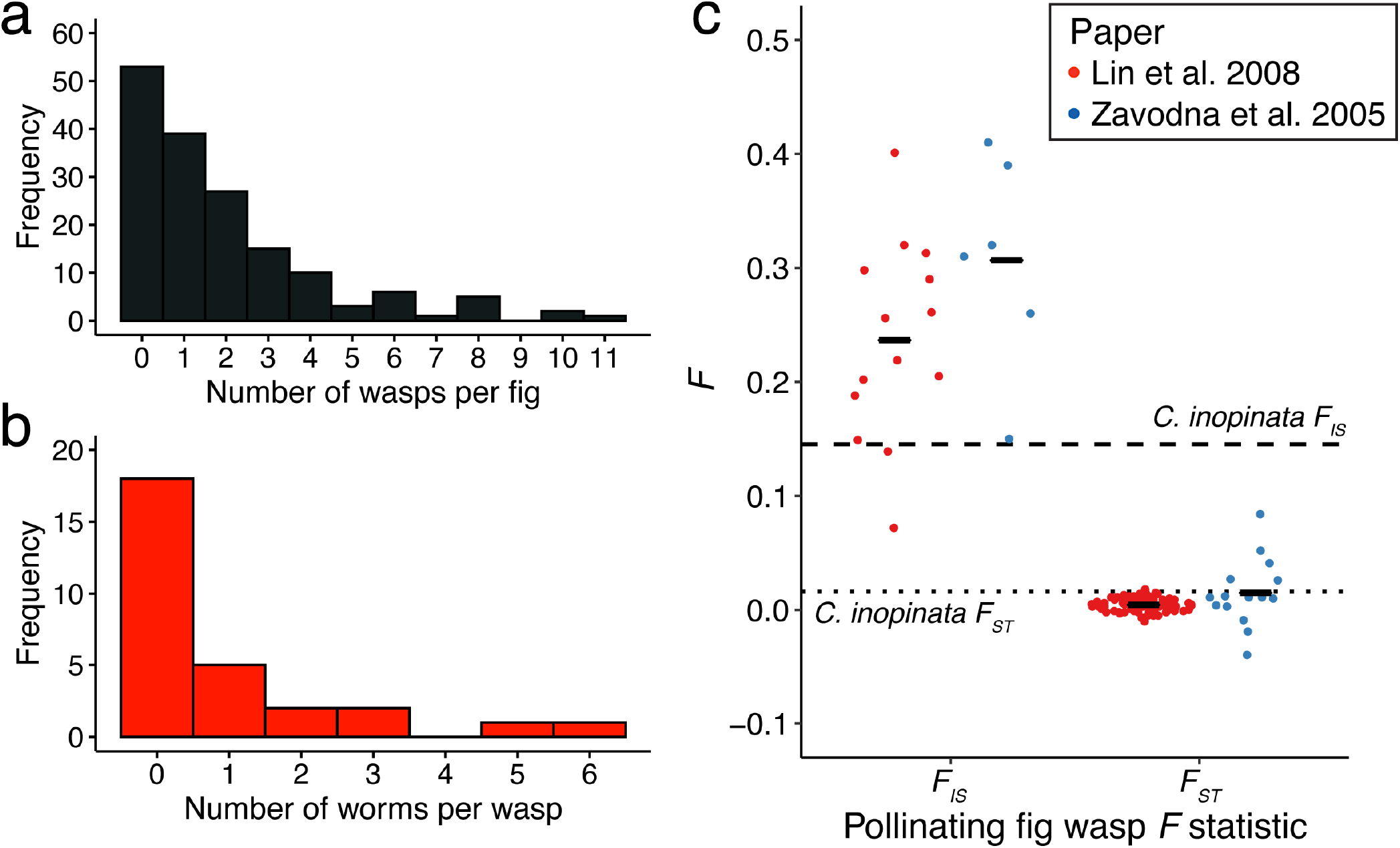
Low *C. inopinata* founding populations in figs and the fig wasp vector of *C. inopinata* has high dispersal and inbreeding. a) The distribution of founding *Ceratosolen bisculatus* pollinating fig wasps among pollinated *F. septica* figs. b) The distribution of *C. inopinata* worms observed traveling on *Ceratosolen* pollinating fig wasps. Data in (a) and (b) are from Woodruff and Phillips *BMC Ecology* 2018 [18]. c) The distribution of *C. bisculatus F*_*IS*_ and *F*_*ST*_ statistics from two previous studies (Lin et al. *Mol. Ecol*. 2008 [20] and Zavodna et al. *J. Evol. Biol*. 2005 [19]). For [20]: 1,502 bp of the COI locus plus 15 microsatellites; 15 localities across Taiwan and Lanyu Island used as populations; 86 trees and 219 (COI) or 398 (microsatellites) wasps; *F*_*IS*_ data from Table 5 and pairwise *F*_*ST*_ data from Table 6. For [19]: five microsatellites; six localities including four islands across Indonesia used as populations; 74 wasps; *F*_*IS*_ data from Table 3 and pairwise *F*_*ST*_ data from Table 5. Both studies revealed low differentiation among fig wasps on islands separated by open ocean ([20], 60 km; [19], 40 km). Black horizontal bars represent means. Dotted and dashed lines denote mean island population pairwise *F*_*ST*_ and mean *F*_*IS*_ respectively among 10kb genomic windows in *C. inopinata* (Figure 3).

### The alignment of genetic diversity among community members

How do these patterns of genetic diversity align across species separated by hundreds of millions of years of evolutionary divergence? Their shared modes of dispersal and small founding populations together drive the synchronization of genomic differentiation in these species. Fig wasps have an atypical life cycle defined by their symbiotic relationship with figs—every generation, female wasps emerge from figs to transfer pollen to a new fig, lay their eggs in fig ovules, and die [10, 16]. The life cycle of *C. inopinata* is also intertwined with the fig-fig wasp symbiosis as *C. inopinata* reproduces in the lumen of young figs, times key developmental decisions to wasp emergence, and disperses to new figs on fig wasps [17, 18]. Thus, as *C. inopinata* and *C. bisculatus* travel together to new resource patches, it is then natural that they would share comparable patterns of *F*_*ST*_ (Fig. 4C). And as fig wasps have been repeatedly demonstrated to be capable of migrating on the wind for tens to hundreds of kilometers (even across spans of open ocean) [19, 20, 29], this is also consistent with low differentiation among island populations in both species. However, as signatures of population differentiation of *F. septica* and its pollinators have been observed across greater geographical distances (including the Philippines, Taiwan, and Okinawa) [27], it is possible that differentiation among *C. inopinata* populations may be found across larger spatial scales. At the same time, founding populations (foundress wasps per fig and nematodes per wasp) are low in both species (Fig. 4A-B), which would lead to mating among close relatives and higher inbreeding coefficients in both species (Fig. 4C). Why exactly such founding populations are so low in both species is unclear. Figs have long been thought to dominate the fig-fig wasp relationship, as pollinating wasps paradoxically pollinate female figs where they are incapable of laying eggs [30]. Figs are likely also advantaged by low foundress wasp numbers, as only one pollinator is required to pollinate the whole fig [30] and would presumably allow for more seed production. At the same time, high nematode loads could impair wasp fitness and dispersal [31, 32], likely contributing to low founding numbers in *C. inopinata* as well. Thus, shared modes of dispersal coupled with low founding numbers constrained by a shared ecosystem drive the alignment of genetic diversity across these divergent species.

### C. inopinata *diversity in its phylogenetic context*

The *Caenorhabditis* genus is noteworthy in the range of intraspecific polymorphism among its constituent species [23-25, 33-37] (Supplemental Figure 2). This is driven largely by the independent evolution of self-fertile hermaphroditism in three species (including *C. elegans*) that have lower levels of polymorphism due to high selfing rates. Conversely, most *Caenorhabditis* species (including *C. inopinata*) are obligate outcrossers [17, 38, 39]. A number of these species appear to harbor exceptional levels of diversity (i.e., are “hyperdiverse” with π ≥ 0.05) suggestive of massive population sizes [40]. *C. inopinata* harbors low levels of polymorphism compared to its hyperdiverse outcrossing relatives while being far more diverse than self-fertilizing *Caenorhabditis* species (Supplemental Figure 2). Its modest diversity as a female/male species may be explained by its vector specialist lifestyle. It is thought that specialists should have lower population sizes and more differentiation among populations compared to generalists because there are fewer opportunities for them to thrive and reproduce in space [37]. Notably, *C. japonica* is another vector specialist that also has lower levels of diversity among outcrossing *Caenorhabditis* [37]. These comparatively lower levels of diversity in *C. inopinata* and *C. japonica* provide evidence for this specialist-generalist variation hypothesis (SGVH; [37]). However, the lack of population differentiation in *C. inopinata* is inconsistent with the expectations of population fragmentation under the SGVH (Fig. 3). This incongruity could result from the notable migration distances of fig wasps [29], the limited spatial area of our study, or both. Moreover, there are more known *Caenorhabditis* vector specialists and generalists whose diversity has not been determined [39]. Exploring diversity in more *Caenorhabditis* species and situating them in their phylogenetic context, as well as the geographic range and extent of population differentiation in *C. inopinata*, will be needed to show if the SGVH holds in this group.

*C. elegans* [22, 23], *C. briggsae* [24], and *C. tropicalis* [25] are notable in that they have elevated levels of diversity on chromosome arms relative to centers. This is consistent with previous observations showing that polymorphism covaries with recombination rate [41]. Diversity in chromosome arms is elevated in *C. inopinata*, but its extent is much lower than that observed in selfing species (Fig. 2B). It is possible that increased effective recombination in obligate outcrossers may blunt the impact of background selection that removes diversity in the chromosome centers of selfing species [23]. An alternative explanation of *C. inopinata*’s flattened genomic landscape of diversity could be its transposon-riddled genome [17], which also co-occurs with a more uniform distribution of protein-coding genes across chromosomes [42]. Further population genomic studies in *Caenorhabditis* outcrossers will be needed to determine if *C. inopinata*’s genomic landscape of polymorphism is representative or an outlier with respect to its mode of reproduction.

### Conclusions

Understanding the causes of spatial genetic structure is a fundamental goal of molecular ecology. Here, we have shown that fig nematodes and their fig wasp vectors share similar patterns of genetic differentiation due to their shared modes of dispersal and their common symbiotic relationship with figs. Interspecific relationships that involve the colocalization and comigration in space of individuals from different species are ubiquitous. Host-pathogen, host-parasite, and phoretic relationships (among others) abound and could lead to similar alignments of genetic diversity among divergent species. Thus, ecological interactions are a potentially widespread driver of genetic structure despite the vast range of geographic distances and body sizes of the species involved.

## Supplemental Information

Supplemental information can be found online at *Journal*.

## Acknowledgements

We thank Anastasia Teterina for offering helpful feedback throughout the development of this work and the preparation of this manuscript. Bill Cresko, Peter Ralph, Andy Kern, and their laboratory members likewise provided helpful comments throughout the development of this work. We thank Waldir Berbel-Filho and Kimberly Moser for providing helpful comments on earlier versions of this manuscript. We thank Tyler Hether for sharing Flip2BeRAD and assisting the processing of bestRAD reads. We thank Erik Andersen and the *C. elegans* Natural Diversity Resource for sharing *C. elegans* variant data. We thank the University of Oregon Genomics and Cell Characterization Core Facility (GC3F) for assistance with Illumina sequencing. This work also benefited from access to the University of Oregon high performance computer, Talapas. This work was supported by funding from the National Institutes of Health to GCW (Grant No. 5F32GM115209-03) and to PCP (Grant Nos. R01GM102511, R01AG049396, and R35GM131838).

## Author Contributions

GCW and PCP devised and designed the project. GCW collected nematodes, performed bioinformatic analyses, and wrote the first draft of the paper; JHW prepared DNA libraries for sequencing; GCW and PCP revised and prepared the final manuscript.

## Methods

### Nematode isolation, sample preparation and sequencing

Animals were collected from the field in previously described work [18, 43]. Briefly, twenty-four individual *C. inopinata* animals were isolated from fresh, dissected *Ficus septica* figs from three Okinawan islands in May 2016. Live animals were fixed in 100% ethanol and kept at –20°C for 3–11 months. Fixed animals were then washed three times in PBS, and individual worms were transferred to individual tubes. Animals were digested with Proteinase K in 20 μl reactions. The proteinase was heat-inactivated (10 min at 95°C), and then half of the reaction was used for linear amplification with the Illustra GenomiPhi V3 amplification kit (GE Lifesciences). DNA was then purified with the Zymo Genomic DNA Clean and Concentrator kit. EcoRI bestRAD libraries were prepared [21], and paired-end 150 bp reads were generated with the Illumina Hi-Seq 4000.

### Genotyping and inference of population genetic statistics

Reads were re-oriented for processing with *Flip2BeRad* (https://github.com/tylerhether/Flip2BeRAD), and the first two base pairs were removed from all reads with *fastx_trimmer* (version 0.0.13; options -f 3 -Q 33; http://hannonlab.cshl.edu/fastx_toolkit/) for downstream processing. Reads were then demultiplexed with *stacks process_radtags* (version 2.0; options -e ecoRI -r -c -q) [44]. Reads were aligned to the reference *C. inopinata* genome assembly [17] with *gsnap* (version 2018-03-25; options --trim-mismatch-score=0 --trim- indel-score=0 --format=sam) [45]. Unique alignments were then extracted (see align_genotype_pop_gen.sh; all data and code associated with this work have been deposited in Github https://github.com/gcwoodruff/inopinata_population_genomics_2020).

Genotypes were called with *bcftools mpileup call* (version 1.9; options *mpileup* -Ou; options *call* -c -Ob – ploidy-file –samples-file) [46]; X chromosome sites in males were called as haploid while female X chromosome and all autosome sites were called as diploid. Alignment files of all samples were then merged with *bcftools merge* (version 1.9; options --info-rules DP:join, MQ0F:join, AF1:join, AC1:join, DP4:join, MQ:join, FQ:join). Sites with coverage <15x and with <80% of samples having genotype calls were removed with *bedtools view* (version 1.9; for coverage: options -i ‘DP>=15’; for missing genotype calls on autosomes: options -i ‘COUNT(GT=“mis”)<5’; for missing genotype calls on X chromosomes: options -i ‘COUNT(GT=“mis”)<4’). Biallelic sites were extracted with *bedtools view* (version 1.9; options: -m2 -M2 -v snps --min-ac 2:minor) and combined with invariant sites to produce a single VCF for inference of population genetics statistics. Additionally, separate VCF files were generated for estimating population genetics statistics of the X and autosomes.

VCF files were processed with *popgenwindows*.*py* (https://github.com/simonhmartin/genomics_general/blob/master/popgenWindows.py) to estimate π (retrieved May 28, 2019; options --windType coordinate -w 10000 -s 10000 -m 160 -f phased) and *F*_*ST*_ (options --windType coordinate -w 10000 -s 10000 -m 160 --analysis popPairDist -f phased -p ishigaki -p iriomote -p yonaguni --popsFile) in 10kb genomic windows. *Stacks populations* [44] was used to estimate site *F*_*IS*_ (version 2.2; options --sigma 3333 --genepop --structure --phylip) and *bedtools map* (version 2.25.0; -o mean -c 4) [47] was used to initially determine the mean *F*_*IS*_ in 50 bp genomic windows. This was done to account for the high number of missing sites resulting from reduced-representation RAD sequencing. After the removal of windows composed only of missing sites, *bedtools map* (version 2.25.0; -o mean -c 4) was used to find mean *F*_*IS*_ in 10kb genomic windows. For comparing π on chromosomes arms and centers between *C. inopinata* and *C. elegans*, windows were normalized by chromosome position by setting the median chromosome base pair to 0 and the end chromosome base pairs to 0.5 (as in [42]). Chromosome “centers” were defined as those genomic windows with normalized chromosomal position < 0.25; chromosome “arms” as those with normalized chromosomal position ≥ 0.25.

*F* statistics data in *Ceratosolen bisculatus* were collected from [20] (Tables 5 and 6) and [19] (Tables 3 and 5). Natural history data for *C. inopinata* occupancy on fig wasps and fig wasp foundress number in *F. septica* figs were communicated in Figures 5, 6, and S1 of [18]. All of these data have been deposited on Github (https://github.com/gcwoodruff/inopinata_population_genomics_2020).

For estimates in *C. elegans*, alignment files (BAM) of twenty-four previously whole genome sequenced *C. elegans* strains (BRC20067, CB4856, CX11271, CX11276, CX11285, CX11314, DL200, DL226, DL238, ECA246, ECA251, ECA36, ED3017, ED3040, ED3048, ED3049, EG4725, JT11398, JU258, JU775, LKC34, MY16, MY23, and N2) were retrieved from the CeNDR database [22] (retrieved June 2021; https://www.elegansvariation.org/data/release/latest). These strains were chosen because they included the “divergent set” of twelve strains (CB4856, CX11314, DL238, ED3017, EG4725, JT11398, JU258, JU775, LKC34, MY16, MY23, and N2; https://www.elegansvariation.org/strains/catalog) plus an additional arbitrarily chosen twelve strains to produce a dataset with a sample size comparable to our *C. inopinata* data. The positions of EcoRI cut sites in the *C. elegans* reference genome were identified with *EMBOSS fuzznuc* (version 6.6.0, -pattern ‘GAATTC’ -complement -rformat gff) [48]. *C. elegans* alignments located within 332 bp of an EcoRI site were extracted with *samtools view* (options -b -L) [49]. Alignments were then genotyped with *bcftools mpileup call* and processed as above.

All analyses and figures for this paper were generated in the R statistical programming language [50]. The R packages “ggplot2” [51], “lemon” [52], “ggforce” [53], “reshape2”[54], and “patchwork” [55] were used.

FASTQ and BAM files have been submitted to the NCBI Sequence Read Archive (SRA; http://www.ncbi.nlm.nih.gov/sra) under accession number PRJNA769443. VCF files have been deposited in Figshare (https://figshare.com/projects/C_inopinata_population_genomics_2021/123973). Sample metadata can be found in supplemental_tables.xls Sheet 1. All other data and code affiliated with this work have been deposited in Github (https://github.com/gcwoodruff/inopinata_population_genomics_2020).

