## Supplemental Figures for "Alignment of genetic differentiation across trophic levels in a fig community"

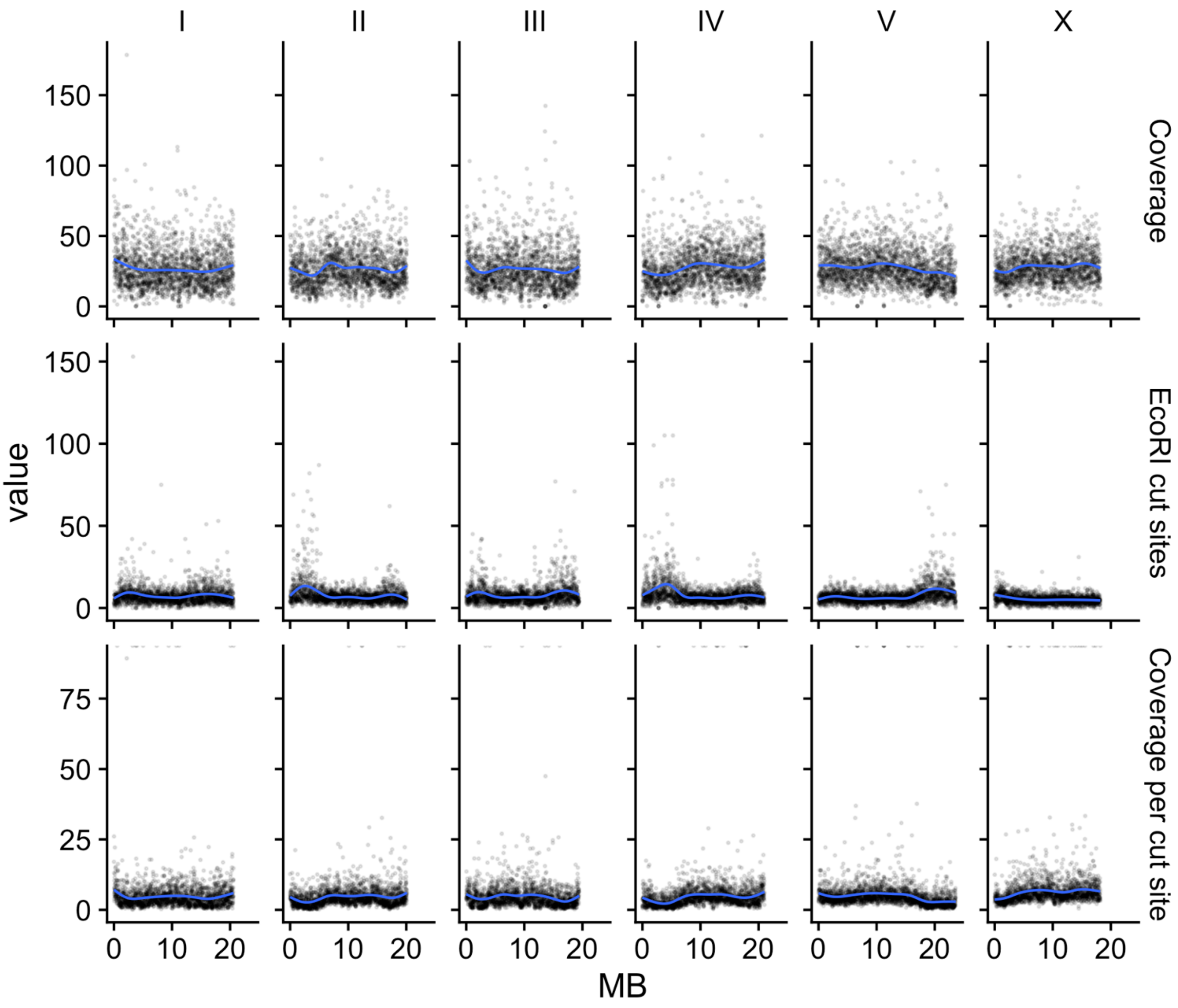


Supplemental Figure 1. Coverage and restriction site across chromosomes. Each column represents a *C. inopinata* chromosome. Top row, mean coverage (x) in 10 kb genomic windows before calling genotypes. Middle row, total number of EcoRI restriction sites in 10 kb genomic windows of the *C. inopinata* reference assembly. Bottom row, (mean coverage)/(number of EcoRI restriction sites) in 10 kb genomic windows.

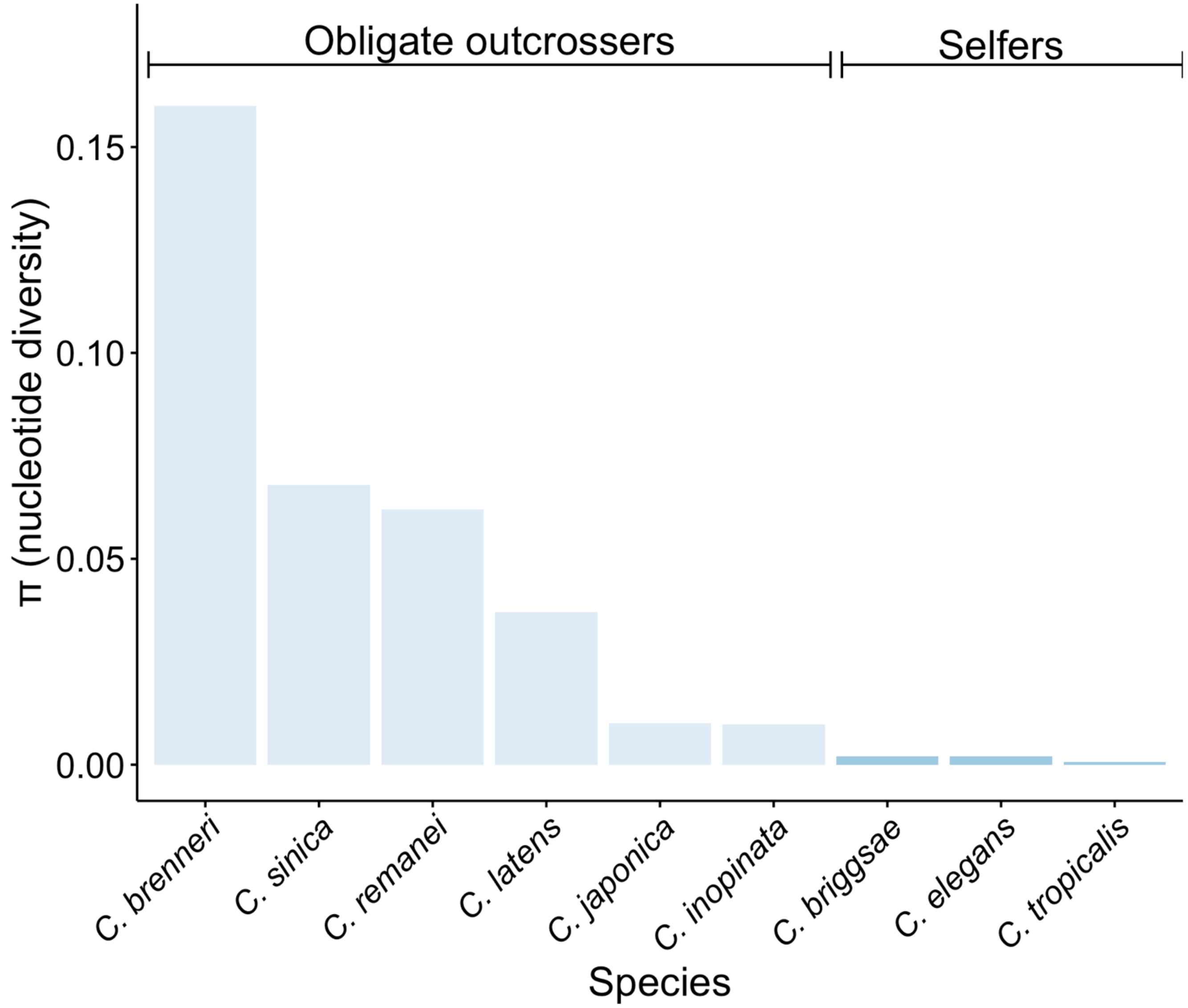


Supplemental Figure 2. Estimates of nucleotide diversity in *Caenorhabtidis* species. Data were retrieved from these sources for each species: *C. brenneri* (0.16; Dey et al. 2013; p. 11056; "we estimate that neutral polymorphism in *C. brenneri* is $\pi_{neu}$ = 16.4%"); *C. sinica* (0.068; Wang et al. 2010; p. 5025; "Nucleotide polymorphism at synonymous sites is very high in our sample of 22 individuals of *C*. sp. 5, averaging 6.84% across seven nuclear loci"); *C. remanei* (0.062; Dey et al. 2012; p. 1260; "with polymorphism in a pooled sample of these populations being somewhat higher (pooled $\pi_{neu}$ = 6.2%...)"); *C. latens* (0.037; Dey et al. 2012; p. 1261; Table 1, Mean $\pi_{neu}$ for *C*. sp. 23 Wuhan, 0.03728); *C. japonica* (0.01; Li et al. 2014; p. 5; "we found an average SNP density at silent sites of $\pi_{si}$ = 0.97%"); *C. inopinata* (0.011, this paper); *C. briggsae* (0.002; Cutter et al. 2006; p. 2024; "Overall silent-site nucleotide polymorphism is low, with $\pi_{si}$ and $\theta_{si}$ estimated to be 0.2%"); *C. elegans* (0.0022, this paper; data from Cook et al. 2017); *C. tropicalis* (0.0006; Gimond et al. 2013; p. 3092; "*C*. sp. 11: $\pi_{si}$ = 0.060%").

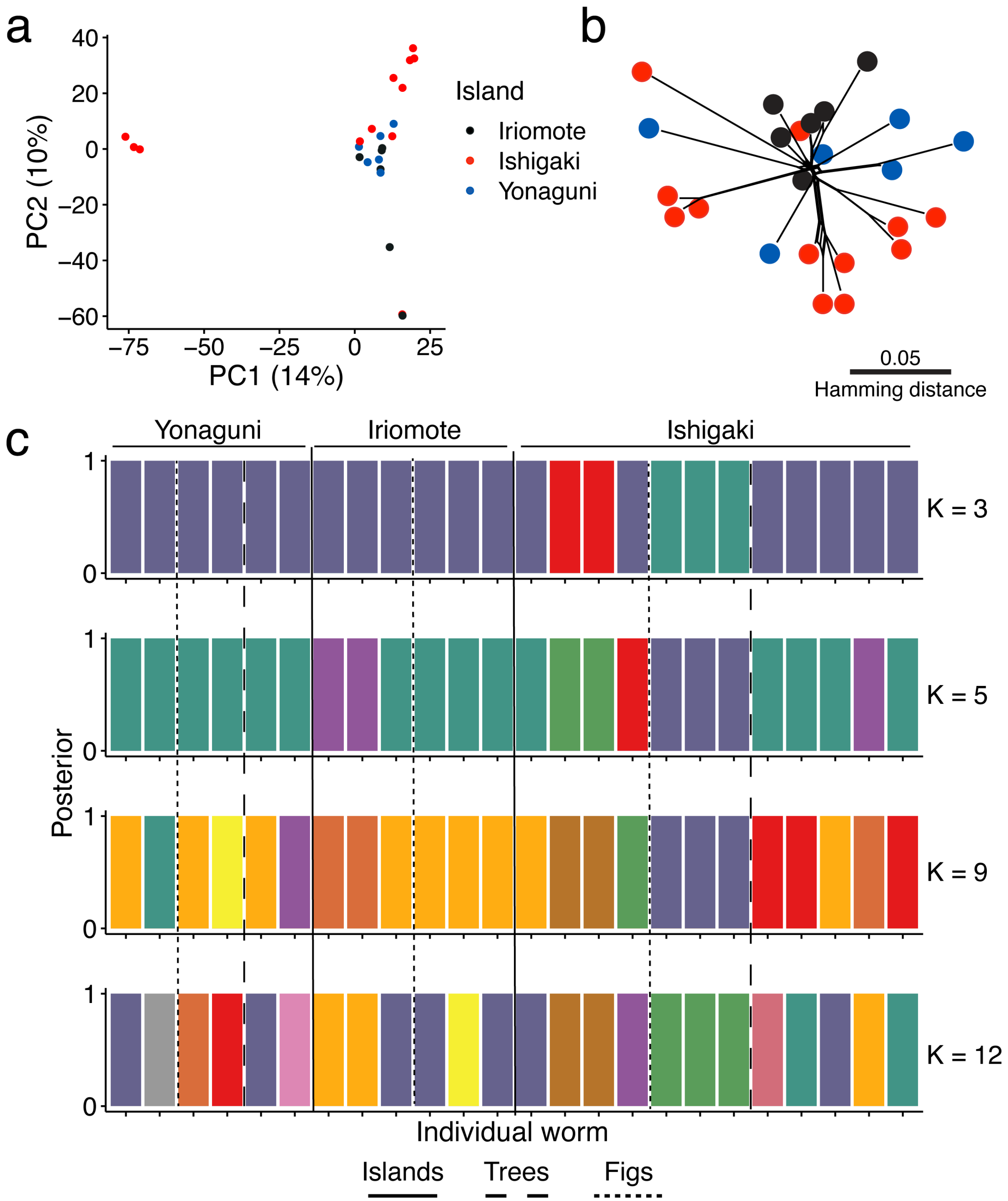


Supplemental Figure 3. *C. inopinata* does not reveal clear genetic structure among island populations. (a) Scatterplot of the first two principal components (PCA) of genetic diversity among individuals. (b) Network distance phylogeny (made with Spitstree). c) Partitioning of individuals into *k* populations by discriminant analysis of principal components. Values of *k* were chosen based on the number of islands (three), trees (five), and figs (nine) animals were sampled from; *k* = 12 had the highest BIC value via *k*-means clustering (Supplemental Figure 4). Color denotes population assignment. Individual worms are ordered by spatial origin; vertical lines separate individuals isolated from the same figs, trees, and islands.

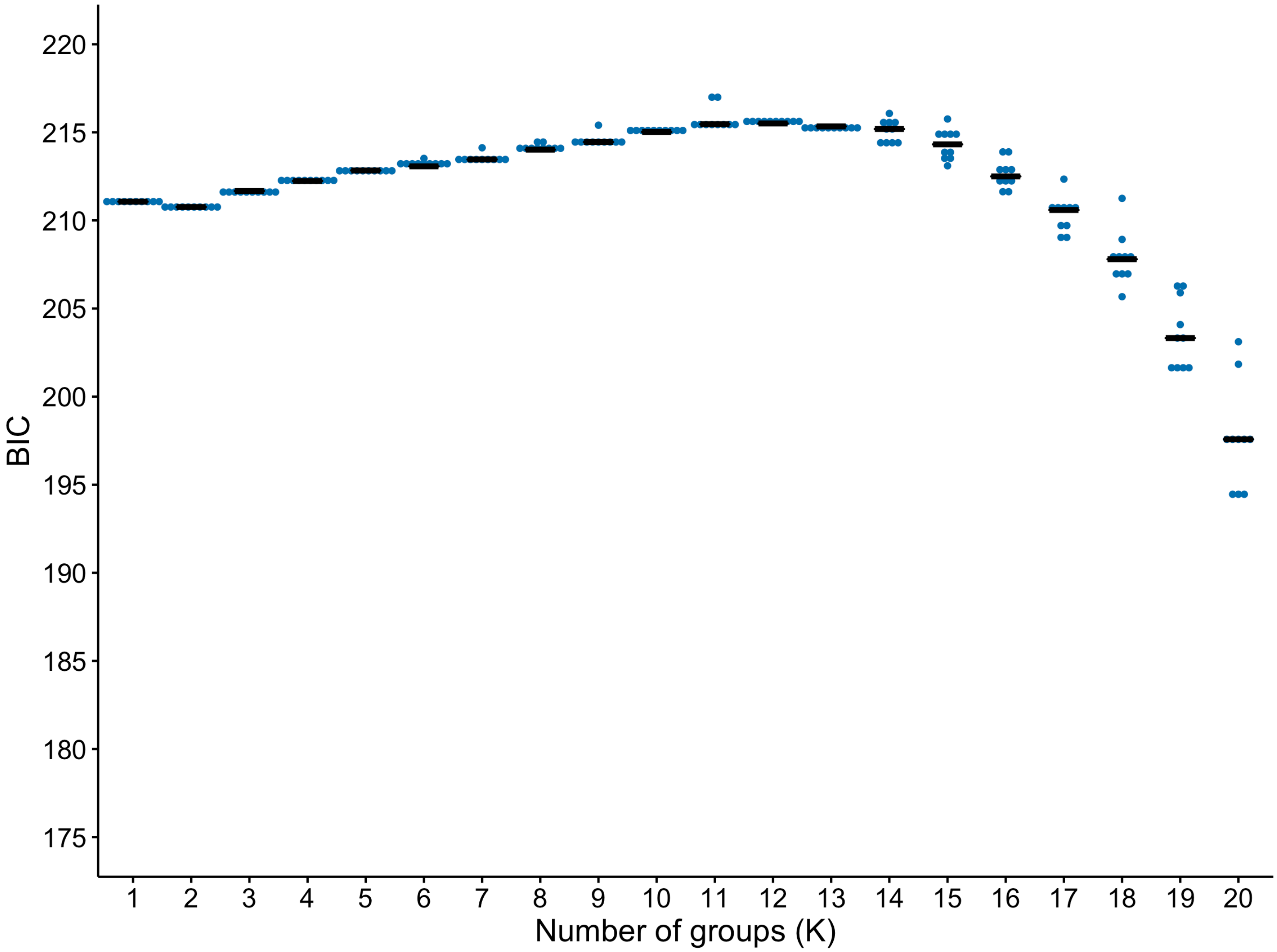


Supplemental Figure 4. BIC for different values of *k* with *k*-means clustering among 24 *C. inopinata* individuals. Clustering was performed 10 times for each value of *k*. Black horizontal bars represent medians.
